# A novel mouse allele of the DNA/RNA helicase senataxin (*Setx^spcar3^*) causing meiotic arrest of spermatocytes and male infertility

**DOI:** 10.1101/2023.04.12.536672

**Authors:** Yasuhiro Fujiwara, Kouhei Saito, Fengyun Sun, Sabrina Petri, Erina Inoue, John Schimenti, Yuki Okada, Mary Ann Handel

## Abstract

An unbiased screen for discovering novel genes for fertility identified the *spcar3, spermatocyte arrest 3,* mutant phenotype. The *spcar3* mutation identified a new allele of the *Setx* gene, encoding senataxin, a DNA/RNA helicase that regulates transcription termination by resolving DNA/RNA hybrid R-loop structures. Although mutations in the human *SETX* gene cause neural disorders, *Setx^spcar3^* mutant mice do not show any apparent neural phenotype, but instead exhibit male infertility and female subfertility. Histology of the *Setx^spcar3^*mutant testes revealed absence of spermatids and mature spermatozoa in the seminiferous tubules. Cytological analysis of chromosome spread preparations of the *Setx^spcar3^* mutant spermatocytes revealed normal synapsis, but aberrant DNA damage in the autosomes, and defective formation of the sex body. Furthermore, *Setx^spcar3^* testicular cells exhibited abnormal accumulation of R-loops compared to wild type testicular cells. Transient expression assays identified regions of the senataxin protein required for sub-nuclear localization. Together, these results not only confirm that senataxin is required for normal meiosis and spermatogenesis but also provide a new resource for determination of its role in maintaining R-loop formation and genome integrity.

## INTRODUCTION

Spermatogenesis produces millions of genetically unique sperm throughout life. Gametes acquire their genetic diversity primarily through meiosis, a special mode of cell division involving exchange of DNA between chromosomal homologs, followed by separation of homologs and chromatids in two divisions. Meiosis is initiated with DNA replication followed by formation of programmed DNA double-strand breaks (DSBs), mediated by the SPO11/TOPO6BL complex [1-4]. During meiotic prophase I, homologous chromosomes (homologs) pair and synapse while undergoing homologous recombination-mediated DSB repair [5-7], resulting in exchange of genetic information, followed by chiasmata formation, the physical manifestation of crossing over. These essential events are mediated by a chromosomal structural framework composed of cohesins and the axial elements (AE) that become the lateral elements (LEs) of synaptonemal complex (SC) [8-10]. These meiotic processes are functionally interdependent and regulated by precise regulation of gene expression and epigenetic modifications [11, 12]. Differential regulation of gene expression is manifest in exceptionally high transcription of autosomal genes in meiotic prophase [13, 14] compared to transcriptional inactivation of the X and Y sex chromosomal genes, which is referred to as meiotic sex chromosome inactivation (MSCI) and is essential for meiotic progression [15-18].

Despite remarkable progress in the past few decades, many of the genes that control these essential meiotic and spermatogenic processes remain unknown and/or understudied for lack of genetic resources. To meet this challenge, we conducted an unbiased phenotype-driven mutagenesis screen using ethyl-nitroso-urea (ENU) [19, 20] to discover novel mutations in genes involved in meiosis and reproduction. Here, we report the phenotype of the *spcar3* (*spermatocyte arrest 3*) mutation identified by ENU mutation. Male *spcar3* mice are infertile, while females are subfertile. Genetic mapping and sequencing identified the *spcar3* mutation as a new allele (*Setx^spcar3^*) of the *Setx* gene, which encodes senataxin, a DNA/RNA helicase. Male *Setx^spcar3^* mutant mice exhibit meiotic arrest with aberrant DNA damage in autosomes and defective sex-chromosome dynamics. By confirming previous analyses of a knockout mutation of *Setx* [21, 22], these results demonstrate the importance of unbiased mutagenesis for producing new alleles to further the study of function of known genes. This newly identified allele of *Setx* not only emphasizes its crucial role in mammalian meiosis and fertility but also provides an additional resource to study structure of the sentaxin protein and determine its role in maintenance of genomic integrity, meiosis, and fertility.

## MATERIAL AND METHODS

### Mice

All mice were obtained from The Jackson Laboratory (JAX) and maintained in research colonies at JAX and The University of Tokyo, Institute for Quantitative Bioscience. The protocols for their care and use were approved by JAX Institutional Animal Care and Use Committee (IACUC) or the Institutional Animal Care and Use Committee at The University of Tokyo (approval #2715, #2809). Mice were euthanized at 8 weeks or 0 to 5 weeks of age to follow the first wave of spermatogenesis. The *spcar3* mouse used in this study was originally produced by the NIH-supported Reproductive Genomics program at JAX [19, 20]. The *spcar3* mutation was induced in a C57BL/6J (hereafter B6) background and subsequently backcrossed to C3H/HeJ mice (hereafter C3H). A *spcar3*-C3H/HeJ congenic line was subsequently generated by backcrossing to C3H over 30 generations. For experimental analyses, homozygous *spcar3* mice were obtained by mating heterozygous *spcar3* mice in the C3H congenic line. All experiments used the congenic *spcar3* line unless indicated. The *spcar3* mice used for experiments were genotyped by PCR amplifying tail DNA with the mismatch primer set. The *Setx^spcar3^* allele in the congenic line was periodically sequenced by a standard Sanger sequencing method for genetic quality assurance. The sequences of the primers used for genotyping are listed in **Table 1**.

**Table 1.**
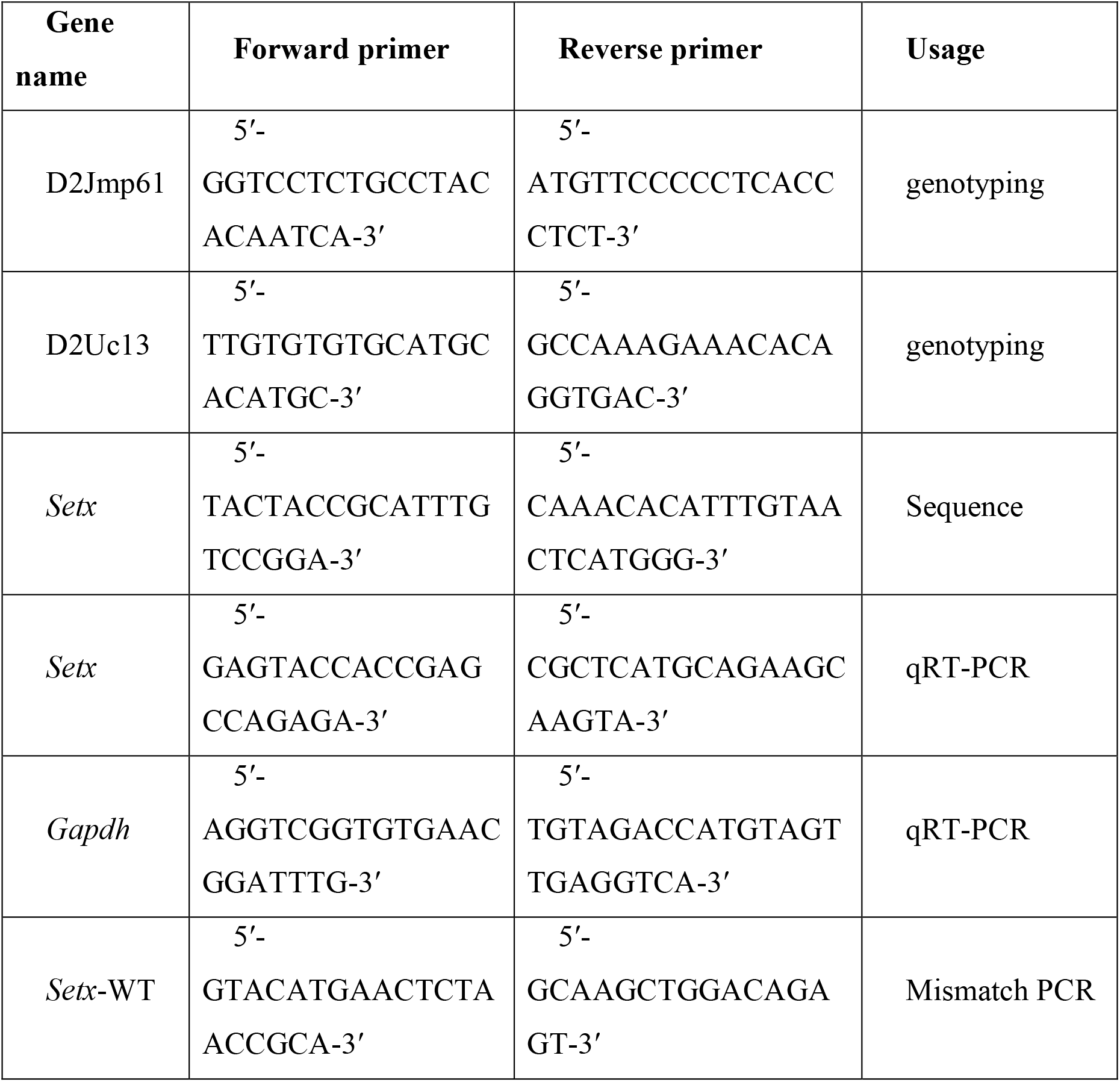
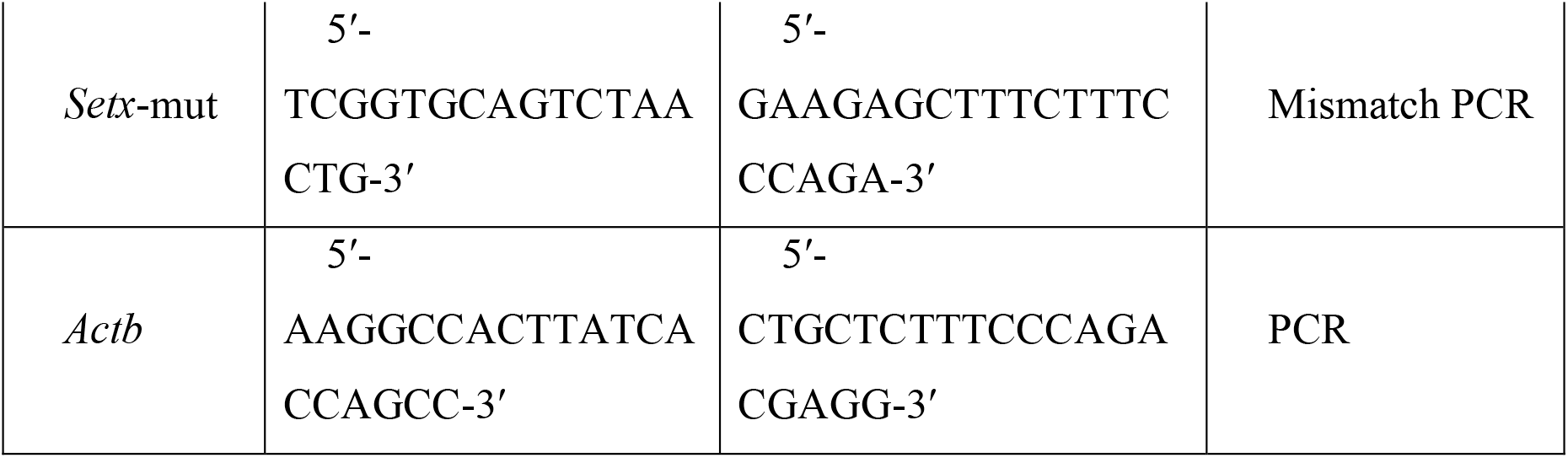
List of primers.

### *Mapping the* spcar3 *mutation*

Adult B6 mice were mutagenized with ENU, and the *spcar3* infertile phenotype was identified in a standard three-generation breeding scheme as previously described [19, 20]. To establish chromosomal linkage of the *spcar3* gene mutation, genome scans using polymorphic satellite makers were performed on DNA obtained from affected and unaffected mice. The offspring were sacrificed for phenotype by histology and genotyped using additional polymorphic microsatellite markers to narrow the candidate region.

For genetic fine mapping, mutant females were crossed to CAST/EiJ (CAST) males, and F2 individuals were sacrificed for phenotype by histology and genotyped with additional polymorphic markers. Mice used for experiments were genotyped by PCR amplifying tail DNA initially with the polymorphic markers D2Uc13 and D2Jmp61 and later with the mismatch primers.

### RNA-seq transcriptome analysis

Testis samples whole testis samples from WT and *spcar3* mice at 18 dpp (four biological replicates) were immersed in RNAlater (R0901, Sigma-Aldrich) until use, then resuspended in QIAzol Lysis Reagent according to the manufacturer’s instruction. Total RNAs were extracted from homogenized samples using QIAGEN RNeasy MiniKit (74104, QIAGEN). The quality of the isolated RNA was assessed using an Agilent 2100 Bioanalyzer instrument (Agilent Technologies, Santa Clara, CA, USA) and RNA 6000 Nano LabChip assay (5067-1511, Agilent Technologies). The mRNA libraries were purified from total RNA using biotin tagged poly dT oligonucleotides and streptavidin coated magnetic beads. The mRNAs were then fragmented and double-stranded cDNA was generated by random priming. The libraries were analyzed for quality using an Agilent 2100 Bioanalyzer instrument (Agilent Technologies) and DNA 1000 LabChip assay (5067-1504, Agilent Technologies). The amplified libraries were sequenced to generate short 100-bp pair-end reads using an Illumina HiSeq 2500 system (Illumina, San Diego, SA, USA) according to the manufacturer’s instructions.

### Accession number

The Gene Expression Omnibus (GEO) accession numbers for the RNA-seq data reported in this paper is GSE224875.

### Mapping and quantification

The sequence reads were trimmed to remove low quality score reads with trimmomatic [23], and the quality of reads was examined using FastQC [24]. Paired-end reads were mapped to the mouse reference sequence (mm10) with the STAR, version 2.7.3a [25]. Binary files were made using SAMtools, version 1.10 [26]. Quantification of gene expression was analyzed using RSEM [27]. Using the bam files, indels in the candidate genes were analyzed using IGV genome browser, version 2.13.0.

### scRNA-seq transcriptome analysis

Single-cell RNA-seq data of young mouse testis was obtained from a previously published report (E-code E-MTAB-6946)[28]. Sequence reads from 5-25 dpp WT mouse testis were aligned to the reference mouse genome data (mm10) using the 10x Genomics Cell Ranger *count* pipeline, version 6.0.2.[29] with the default setting, and the multiplexed samples were aggregated using the Cell Ranger *aggr* pipeline with the default setting. Cell populations were clustered into seven groups by k-means clustering method and plotted by uniform manifold approximation and projection (UMAP) using 10x Genomics Loupe Browser software version 6.2. According to the previously reported method [30], cell types or spermatogenic developmental stages of clustered populations were identified based on the expression pattern of known stage-specific marker genes, including *Stra8* for preleptotene spermatocytes *Sycp3*, *Piwil1*, *Spo11*, and *Dmc1* for meiotic spermatocytes, *Rec8* for spermatocytes and round spermatids, *Sox9* and *Wt1* for Sertoli cells, *Cyp11a1*, *Hsd3b1*, and *Insl3* for Leydig cells (data not shown).

### Histology

Testes were fixed in Bouin’s solution and paraffin-embedded. Sections (5 μm) were stained with Periodic Acid Schiff (PAS) by standard procedures. The TUNEL assay was performed using the *in situ* Cell Death Detection Kit (11684817910, Roche Applied Science, Penzberg, Germany) according to the manufacturer’s instructions. Bright-field images were acquired using a Leica Leitz DMRXE upright microscope equipped with a DCF 300FXRI camera and Leica FireCam software (Leica Microsystems, Bannockburn, IL). Fluorescent images were acquired using a Leica SP5 confocal microscope (Leica Microsystems, Wetzlar, Germany).

### R-loop detection

Testes were fixed in 4% PFA at 4 ℃ overnight and embedded with O.C.T. Compound (Sakura Finetek Japan, Tokyo, Japan). Frozen sections (10 μm) were boiled for 1 min in 0.01 M sodium citrate (pH 6.0) for antigen retrieval. For an R-loop negative control the preparation was treated with 5 U RNaseH (M0297S, NEB Japan, Japan) in 1× RNaseH Reaction Buffer (NEB Japan) at 37 ℃ overnight. The treated samples were incubated with the S9.6 primary antibody (**Table 2**) at 4 ℃ overnight. Corresponding secondary antibodies conjugated with Alexa Fluor 488, 594, and 647 (Molecular Probes, Invitrogen, Carlsbad, CA, USA) were used at 1:500 dilution at RT for 1 h. DNA was counterstained with DAPI (VECTASHIELD H-1200, VECTOR Laboratories, CA, USA). Images were captured with a DeltaVision Elite microscope equipped with an sCMOS camera, and images for spread samples were deconvoluted and stacked using a DeltaVision SoftWorx software (Cytiva, Massachusetts, USA).

**Table 2.** List of antibodies.

| Antigen | Company | Cat number | Host | Dilution |
| --- | --- | --- | --- | --- |
| SETX-C | This study | n.a. | Rabbit | 1:250 (IF),<br>1:100(simple<br>western) |
| S9.6 | Sigma-Aldrich | MABE1095 | Mouse | 1:250(IF) |
| DMC1 | Santa Cruz | sc-8973 | Goat | 1:50 |
| HA(12CA5) | Roche | 14702500 | Mouse |  |
| P-H2AFX | Millipore | 05-636 | Mouse | 1:200 |
| HIST1H1T | Handel lab | Cobb et al.1999 | Guinea<br>pig | 1:500 |
| MLH1 | BD pharmingen | 51-1327GR | Mouse | 1:50 |
| RAD21L | Watanabe lab | Ishiguro et al.<br>2011 | Rabbit | 1:500 |
| RAD51 | Santa Cruz | sc-8349 | Rabbit | 1:100 |
| SPATA22 | Protein tech | 16989-1-AP | Rabbit | 1:100 |
| SYCP1 | Santa Cruz | sc-20837 | Goat | 1:100 |
| SYCP3 | Handel lab | Eaker et al. 2001 | Rat | 1:1000 |
| TEX12 | ProteinGroup | 17068-1-AP | Rabbit | 1:100 |
| HPRT1 | Proteintech | 15059-1-AP | Rabbit | 1:50 |
| Alexa Flour<br>488 donkey<br>anti-rabbit | Invitrogen | A11008 | Donkey | 1:1000 |
| Alexa Flour<br>488 donkey<br>anti-mouse | Invitrogen | A11029 | Donkey | 1:1000 |
| Alexa Flour<br>555 goat anti-<br>rat | Invitrogen | A21434 | Goat | 1:1000 |
| Alexa Flour<br>568 goat anti-mouse | Invitrogen | A11004 | Goat | 1:1000 |
| Alexa Flour<br>488 goat anti-<br>guinea pig | Invitrogen | A11073 | Goat | 1:1000 |
| Alexa Flour<br>488 rabbit<br>anti-goat | Invitrogen | A11078 | Rabbit | 1:1000 |
| Alexa Flour<br>647 donkey<br>anti-mouse | Invitrogen | A31571 | Donkey | 1:1000 |

### Exogenous expression of SETX in mouse testis

To generate an expression vector for HA-tagged full-length SETX, total RNA was extracted from mouse testis using the RNeasy extraction kit (74104, Qiagen, Hilden Germany) according to the manufacturer’s instruction. *Setx* cDNA was amplified by RT-PCR. The cDNA was inserted into a pAc vector. The pAc-*Setx*-HA plasmid DNA vector was injected into live-mouse testes based on a previously reported method with some modifications [31, 32]. Briefly, mice at 20 dpp for WT mice and 30 dpp for *spcar3* mice are anesthetized, and 50 µg of the plasmid DNA in HBS buffer (21 mM HEPES, 137 mM NaCl, 5 mM KCl, 0.7 mM Na_2_HPO_4_・12H_2_O, 0.1% Glucose) was injected into the testis using a glass capillary under a stereomicroscope. After 60 min, electric pulses were applied four times at 50V for 50 msec with 50 msec intervals, followed by the same electric pulses in the reverse direction, using an electroporator NEPA21 (NepaGene, Chiba, Japan). The testes were then placed in the abdominal cavity. 24 h after electroporation, the mice were euthanized. The testes were extracted and used for cell preparations as described below.

### Surface-spread chromatin preparations

Spermatocytes were collected by centrifugation, surface-spread in wells of multi-spot Shandon slides (Thermo Fisher Scientific, Waltham, MA, USA), and fixed in 1 % PFA as previously described [33-35]. As previously described, primary and secondary antibodies (**Table 2**) were used [35]. DNA was counterstained with DAPI (VECTASHIELD H-1200, VECTOR Laboratories, CA, USA). Images were acquired with a ZEISS Axio Imager.Z2 equipped with a Zeiss AxioCam MRm CCD camera (ZEISS, Jena, Germany), an SP5 confocal microscope (Leica Microsystems, Wetzlar, Germany), or a DeltaVision Elite microscope equipped with an sCMOS camera and images for spread samples were deconvoluted and stacked using a DeltaVision SoftWorx software (Cytiva, Massachusetts, USA).

### mRNA Extraction and qRT-PCR

Total RNA was extracted from isolated male germ cells or whole testes using the RNeasy extraction kit (74104, Qiagen, Hilden Germany) as described above. Transcript-specific primers were designed to span introns and the *spcar3* mutation site and used in one-step RT-PCR reactions (210210, OneStep RT-PCR kit, Qiagen). Primer sequences for qRT-PCR are listed in **Table 1**. Quantitative RT-PCR (qRT-PCR) was performed on the Applied Biosystems 7500 Real-Time PCR System (Applied Biosystems, Foster City, CA) using the QuantiTect SYBR Green® RT-PCR kit (204243, Qiagen) as previously described [36]. Gene-specific primers (Table 1) were used to determine genes’ overall relative expression levels tested according to the standard curve method. SYBR Green® was used to detect the double-stranded DNA produced during the amplification reaction. Reactions were performed using approximately 10 ng of total RNA. One-step RT-PCR reactions were performed in 25-μl volumes as directed by the manufacturer for 40 cycles. For relative quantification method [37], a gene-specific standard curve was established using aliquots of the same stock of RNA for the experimental sample (total RNA extracted simultaneously from multiple 17-dpp testes). Because the same stock of RNA was used to prepare all standard curves, the relative quantities determined for a given gene using this method could be compared across individual experiments. All reactions were performed in duplicate on the same three independent sets of germ cells, and specificity was assessed by melting curve analysis. qRT-PCR results were normalized to their corresponding *Gapdh* and are presented as the mean normalized expression in 10 ng of RNA. Data are reported as mean ±SD.

### Antibody production

Polyclonal antibody against the C-terminus of mouse SETX-C were generated by immunizing rabbits. Mouse cDNA encoding Setx (aa. 1900-2701) was subcloned into pET15b (Novagen) and transformed to *E.coli* strain BL21-CodonPlus(DE3)-RIPL (Agilent). The expressed recombinant protein was solubilized in a denaturing buffer (6 M HCl-Guanidine, 20 mM Tris-HCl[pH 7.5] and purified by Ni-NTA (30210, QIAGEN) under denaturing conditions. The antibody was affinity-purified from the immunized serum with immobilized antigen peptides on CNBr-activated Sepharose (GE healthcare).

### Immunodetection of proteins by Simple Western blotting

The expression of SETX in the whole testis from WT and *spcar3* mice was assayed by a Simple Western assay [38] using a Jess (ProteinSimple, Santa Clara, CA, USA). Whole testis was homogenized with lysis buffer (20 mM Tris-HCl (pH 7.4), 200 mM KCl, 0.4 mM EDTA, 0.1% Triton X-100, 10% glycerol, 1 mM 2-mercaptoethanol) containing a protease inhibitor cocktail (P8340, Sigma Aldrich, MO, US). Samples were run on a Jess Separation Capillary Cartridge (SM-W004, ProteinSimple). Primary antibodies used were rabbit polyclonal anti-SETX-C and HPRT1 antibodies (**Table 2**). The antibody was diluted in an antibody diluent buffer (042–203, ProteinSimple) that was also used as a blocking buffer. Secondary antibodies used were anti-rabbit IgG antibody (042–206, ProteinSimple). Data were analyzed using the software Compass for SW 4.0 (ProteinSimple).

## RESULTS

### Spcar3 *mutant mice exhibit male infertility with arrest during the pachytene stage of meiosis I*

The male-restricted infertile *spcar3* phenotype (**Figure 1A**) was identified in a phenotype screening for infertility by the Reproductive Genomics Project at The Jackson Laboratory [19, 20]. In contrast to the male infertility, female homozygous *spcar3* mice were sub-fertile with reduced litter size (**Figure 1A**). Male *spcar3* homozygous mice exhibited reduced testicular weight after 21 dpp (days postpartum) (**Figure 1B**), with lack of mature spermatozoa in the lumen of seminiferous tubules. To determine developmental timing of spermatogenic arrest, we analyzed histological sections of testes from young pups **Figure 1C**). At 14 dpp, when in wild type (WT) males the most advanced spermatogenic cells were early pachytene spermatocytes, no significant differences were observed between WT and *spcar3* homozygous mice. By 18 dpp, when meiotic spermatocytes at the late pachytene substage were observed in WT testes, germ cells with darkly condensed chromatin were observed in *spcar3* homozygous mice. At 21 dpp, round spermatids were observed in the WT seminiferous tubules, while no spermatids were observed in testes of *spcar3* homozygous males (**Figure 1C**). Adult male *spcar3* homozygous mice also showed cells exhibiting aberrantly condensed chromatin in the tubules (**Figure 1C**). A TUNEL assay exhibited a notable increase in the frequency of TUNEL-positive cells in the testicular tubules of *spcar3* homozygous mice compared with that of WT mice, indicating apoptotic cell death (**Figure 1D**). To more specifically pinpoint the time of spermatogenic arrest, we immunolabeled spread chromatin of spermatocytes for H1T, a marker for mid-to-late pachytene substages [39]. No spermatocytes in homozygous *spcar3* testes were H1t-positive (**Figure 1E**), indicating that spermatogenic arrest occurs at the early-to-mid pachytene substage of meiosis I in the *spcar3* germ cells.

**Figure 1.**
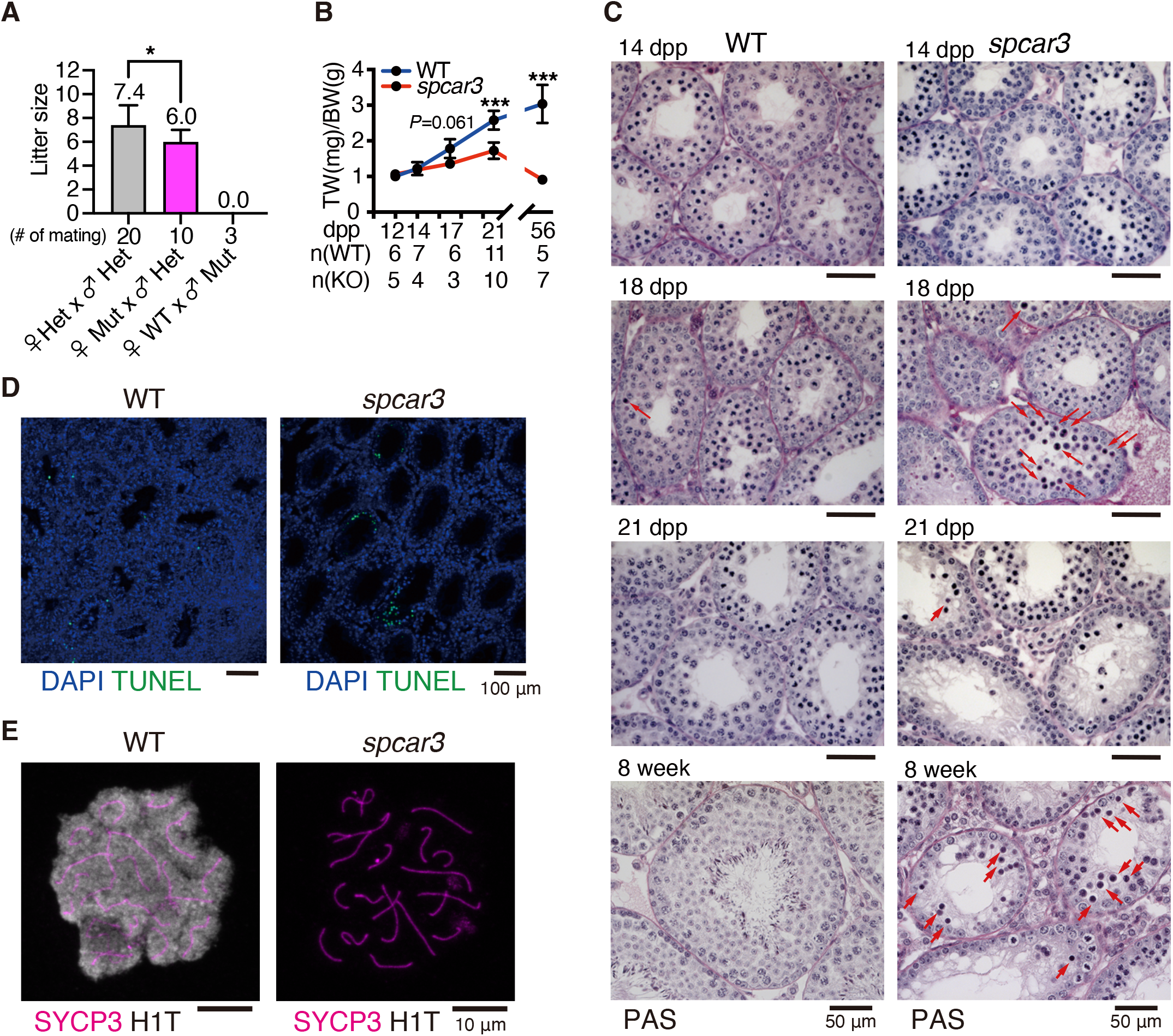
Spermatogenic arrest at the pachytene substage of meiosis I in the *spcar3* mutant mice. **A**. Bar plot showing litter size after mating with pairs indicated on x-axis. Standard deviations are indicated; the statistical significance was determined by non-parametric one-way ANOVA followed by Turkey’s test. * *P* < 0.05. **B.** Line plot showing the developmental trajectory of testis/body weights TW (mg)/BW (g) in WT (blue line) and *spcar3* (red line) mice. Bars indicate standard deviation. Sidak’s multiple comparison test was used to determine statistical significance. *** *P* < 0.001. The numbers below indicate the number of mice examined. **C**. PAS-stained histological sections of testes from WT and the *spcar3* mutant mice providing a developmental series from 14 dpp to 8 weeks, encompassing the first wave of spermatogenesis. Spermatogenic defects in mutant testes were observed after 18 dpp. The red arrows indicate darkly stained degenerating cells. Scale bars are 50 μm. **D**. TUNEL-staining revealed an increased number of apoptotic cells (green) in the testes of *spcar3* mutant mice. DNA is stained by DAPI (blue). **E**. Immunostaining of spread chromatin for the stage-specific marker protein H1T (gray), labeling post-mid-pachytene stages revealed that H1T was expressed in the WT spermatocytes but not in the mutant *spcar3* spermatocytes. SYCP3 (magenta), a component of the synaptonemal complex (SC), was immunolabeled to confirm meiotic stage as pachytene.

### *Normal chromosomal axis formation and synapsis in* spcar3 *mutant spermatocytes*

We further assessed the meiotic progression of the *spcar3* homozygous spermatocytes by immunolabeling spread chromatin of both WT and *spcar3* spermatocytes for axis components, including SYCP3, the main component of axial element (AE), as well as SYCP1 and TEX14, components of central element (CE). The immunolabeling revealed that both WT and *spcar3* spermatocytes exhibited full co-localization of SYCP3 and CE components on their autosome axis (**Figure 2A**), indicating normal synapsis in *spcar3* spermatocytes. We further examined cohesin loading by immunolabeling TEX12 and RAD21L, a component of the meiosis-specific cohesin complex, but no differences were observed between WT and *spcar3* spermatocytes (**Figure 2B and C**). These findings suggest that axis formation and synapsis are normal in mutant *spcar3* mice.

**Figure 2.**
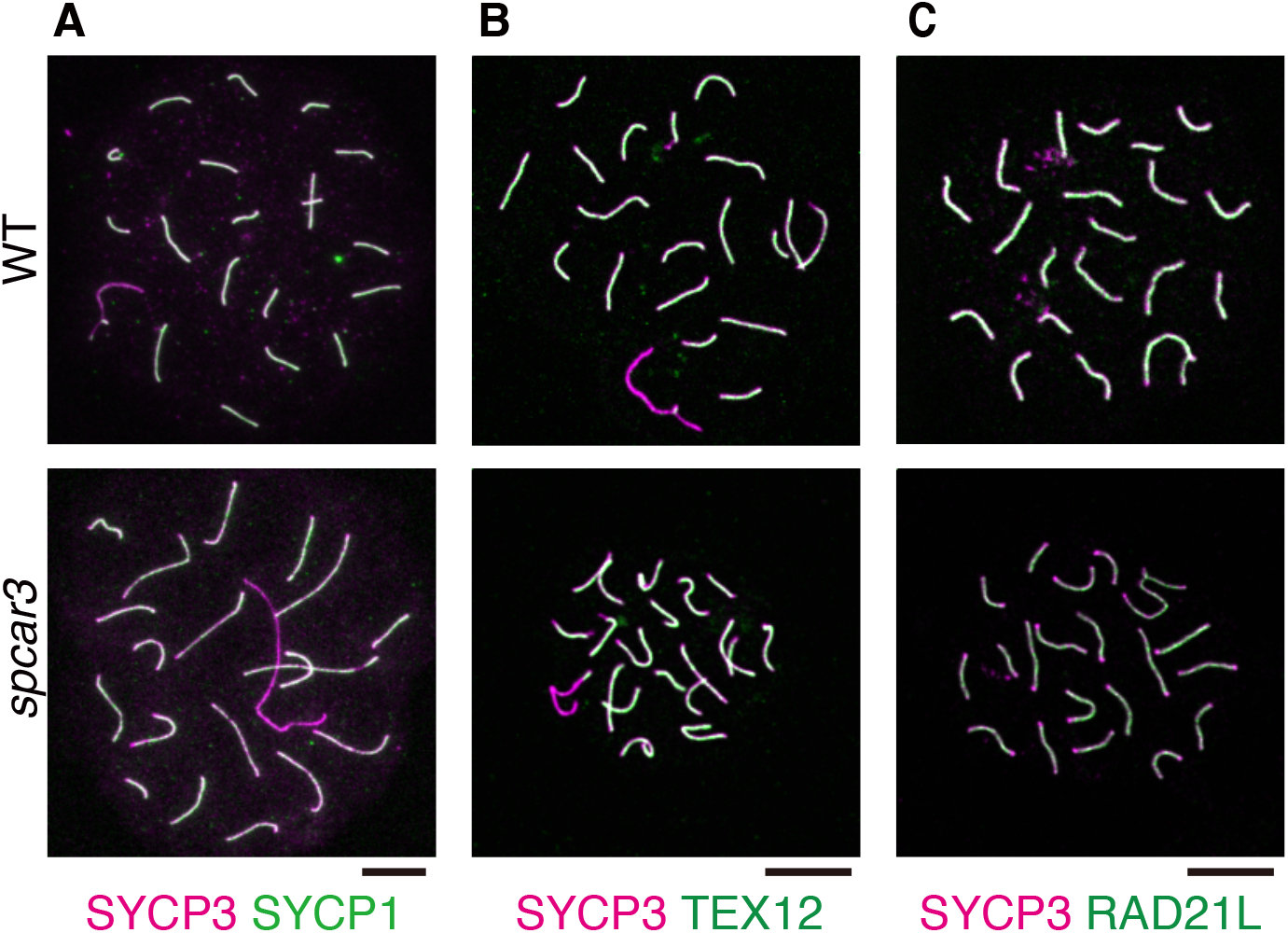
Normal synapsis and axis formation in the mutant *spcar3* spermatocytes. **A-C.** Spread chromatin of spermatocytes from WT and mutant *spcar3* mice were immunolabeled for SYCP1 (A, green) and TEX12 (B, green), components of a central element of the SC, and RAD21L (C in green). SYCP3 (magenta), a lateral element component of SC, was immunolabeled for staging the cell. Scale bars are 10 µm.

### *Spermatocytes of* spcar3 *mutants exhibit aberrant DNA damage repair*

Next, we evaluated DNA damage and its repair machinery by immunolabeling γH2AX, a marker for DNA double-strand breaks (DSBs). DSBs detected by γH2AX were observed only in the XY chromosomes in WT pachytene spermatocytes. In contrast, *spcar3* pachytene spermatocytes exhibited γH2AX in both the XY chromosomes and the autosomes (**Figure 3A**). About 80% of *spcar3* homozygous pachytene spermatocytes, in comparison to only 10% of WT pachytene spermatocytes, showed abnormal autosomal γH2AX localization (**Figure 3A,B**). The majority of autosomal meiotic DSBs in WT spermatocytes are repaired through DNA repair factors DMC1/RAD51 and intermediate mediators, including SPATA22 on the axis [40, 41]. We found that foci of these repair factor decreased from higher than 200 (leptotene/zygotene) to less than 100 by early pachytene (**Figure 3C,D,E,G**) in WT spermatocytes. Although no differences in the numbers of DMC1/RAD51/SPATA22 foci were observed in the *spcar3* spermatocytes during leptotene/zygotene stages, numbers of foci for these proteins were significantly higher than WT at early pachytene (**Figure 3C,D,E,G**). Notably, most of the DMC1/RAD51/SPATA22 foci in the *spcar3* spermatocytes were on the autosomes (**Figure 3C,D,E**), suggesting that the initial formation of DSB generated by SPO11/TOPO6BL and the early steps of DNA damage repair machinery were maintained in the *spcar3* spermatocytes. However, DSBs remained unrepaired on the autosomes of the *spcar3* spermatocytes during the pachytene stage due to either an aberrant repair process during the pachytene stage or an unknown source of *de novo* DSB. Finally, we immunolabled for MLH1, a marker of crossing over that is present in WT spermatocytes at late pachytene [42]. No MLH1 foci were observed in the *spcar3* spermatocytes (**Figure 3F,G**), consistent with findings that DNA damage is not repaired, impeding completion of recombination.

**Figure 3.**
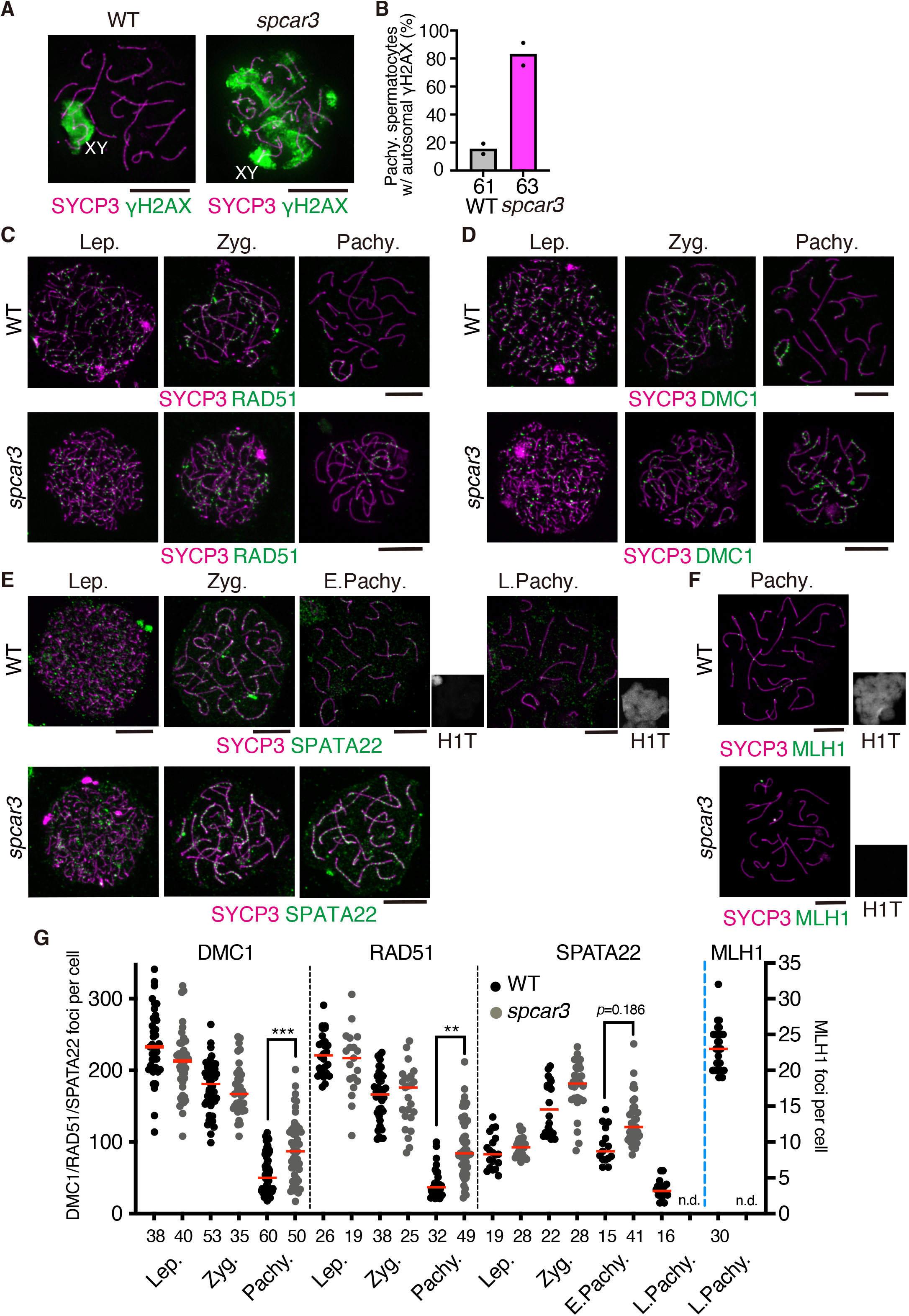
Unrepaired DNA damage in the pachytene-like mutant *spcar3* spermatocytes. **A-B.** Immunolabeling of spread chromatin of spermatocytes from WT and mutant *spcar3* mice for γH2AX (green) to identify the DNA damage sites. SYCP3 (magenta) was immunolabelled to identify meiotic substage. B. Bar graph shows the percentage of pachytene spermatocytes with autosomal γH2AX. The numbers indicated below bars are the number of mice used for quantification. **C-F**, Spread chromatin of spermatocytes from WT and mutant *spcar3* mice were immunolabeled for recombinases RAD51 (C in green) and DMC1 (D, green), the meiosis-specific recombination intermediate SPATA22 (E in green), and the crossover factor MLH1 (F in green). SYCP3 (magenta) and H1T (a marker for post-mid-pachytene germ cells, thumbnail images next to the merged images in E and F in grey) were immunolabeled for staging spermatocytes. G. This dot plot shows quantification for the number of recombination-related foci, with black dots representing WT samples and grey dots representing mutant *spcar3* cells. Only those foci associated with SYCP3 threads were counted. Statistical significance was determined by non-parametric one-way ANOVA followed by Turkey’s test. Red bars represent the median count. \*\**P* < 0.01, \*\*\**P* < 0.001. n.d.: not detected. The number below indicates the number of cells observed. Scale bars are 10 µm.

### *The* spcar3 *mice have a 2 bp deletion mutation in the* Setx *gene*

To identify the causative mutation for the *spcar3* phenotype, we performed linkage mapping analysis using the recombinant F2 mice (**Figure 4A**, see Methods for details). The *spcar3* mutation was mapped to ∼0.4 Mb region of the Chr. 2 between D2Mit83 and D2Mit295 using 15 affected and 30 normal F2 male mice (**Figure 4A**). This region harbors 28 genes. Using the RNA-seq data set of *spcar3* testis at 18 dpp, we examined the mRNA sequences of all 28 candidate genes and identified a two bp-deletion in the *Setx* gene. By amplifying the genomic region of the *Setx* gene in homozygous *spcar3* mice by PCR, we confirmed the two bp-deletion in the *Setx* gene (**Figure 4B**). This two bp-deletion caused a frameshift that resulted in 9 amino acid substitution followed by a premature stop codon at aa 539 (**Figure 4B,C**). Indeed, the *spcar3* mutation abolished the full-length SETX in the testis (**Figure 4D**). Henceforth we refer to the mutant allele as *Setx^spcar3^*. To generate a *Setx^spcar3^* congenic strain, we designed a PCR genotyping protocol using a set of mismatch primers (**Figure 4E,F**). We backcrossed the genotype-confirmed *Setx^spcar3^* allele to the C3H strain for more than 30 generations. The phenotype of the resultant congenic *Setx^spcar3^* strain was similar to that of the *spcar3* founder strain. Notably, this phenotype of male infertility with meiotic arrest and female sub-fertility of the *Setx^spcar3^* allele on a defined (C3H) genetic background reflects that of the previously reported *Setx-*KO allele [22], where the modified allele is on a different background. These results suggest that the phenotype has high expressivity and is not obviously modified by genetic background.

**Figure 4.**
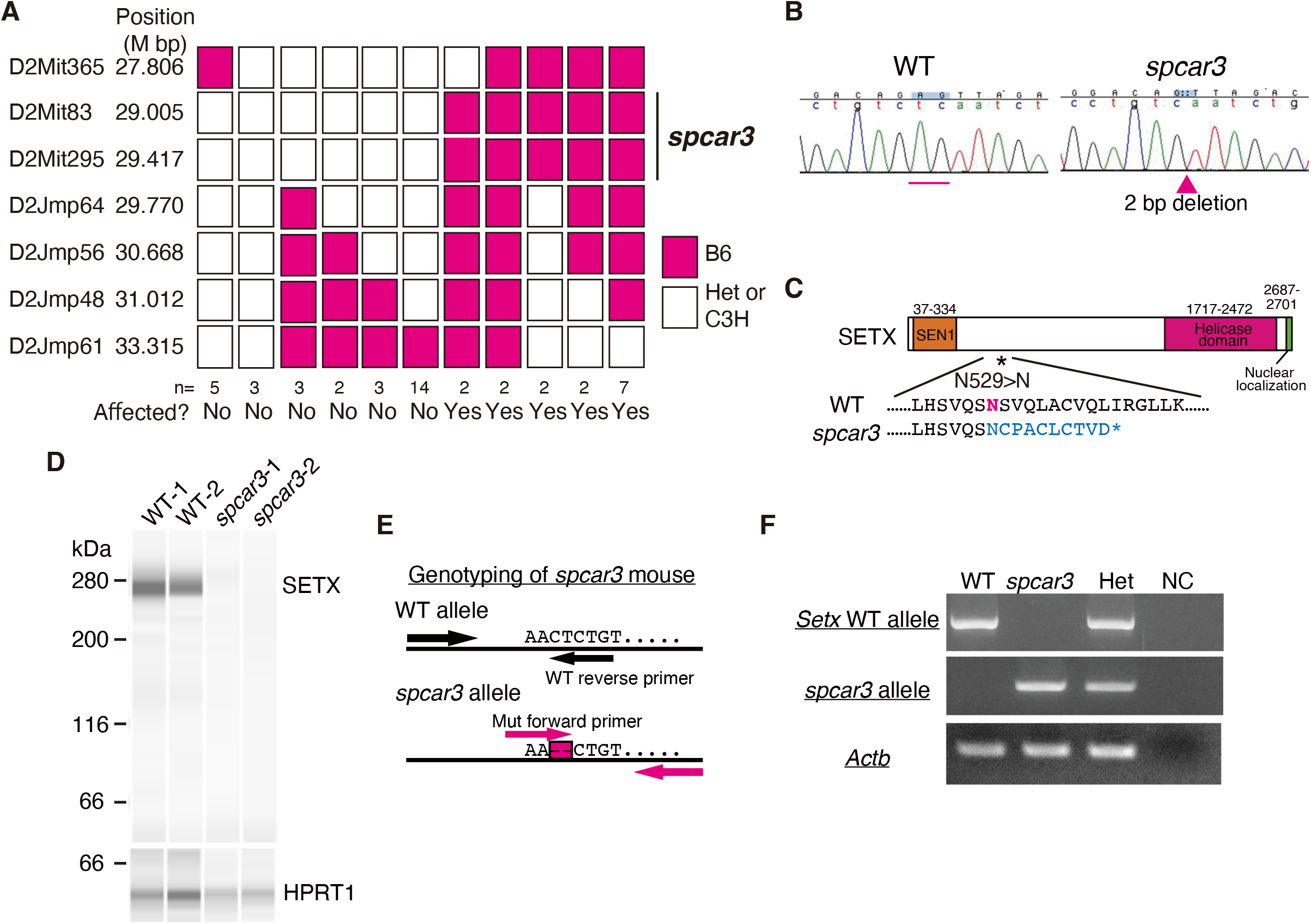
The *spcar3* mutation is in the senataxin (*Setx*) gene. **A**. Fine mapping of the *spcar3* mutation was done by linkage analysis using 7 microsatellite markers for affected and normal F2 mice obtained from a mating between +/*spcar3* and CAST/EiJ mice. Positions of microsatellite markers are indicated in cM according to the mouse genome sequence (NCBI Build 38). The candidate region of the *spcar3* mutation is indicated by the vertical bar on the right. The red and white boxes represent B6 sequence homozygosity and heterozygosity or C3H homozygosity, respectively. The number of mice used for the fine mapping is displayed below the boxes. **B**. The nucleotide sequences of the *Setx* gene in WT and the *spcar3* mice. The site of the *spcar3* mutation containing a 2-bp deletion in the *Setx* is indicated by a red bar in the WT genome and the red arrowhead in the *spcar3* genome. **C.** The two-bp deletion *spcar3* mutation causes frameshift and a premature stop codon, leading to a loss of the C-terminus of SETX protein in *spcar3* mouse. Asterisk indicates the site of the *spcar3* mutation. **D.** Western blot analysis confirmed loss of SETX protein in the *spcar3* mutant testes. HPRT1 is a loading control. **E.** The *spcar3* mutation is genotyped using a mismatch primer set for the WT allele and *spcar3* allele. **F.** Electrophoresis of PCR products using mismatch primers for the *Setx* WT allele and *spcar3* allele and a primer set for *Actb*. NC, negative control.

### Nuclear localization of SETX in meiotic spermatocytes

Since the expression dynamics of SETX during meiotic progression have not been fully elucidated, we assessed expression of *Setx* transcript by quantitative RT-PCR of testes during the first wave of spermatogenesis, in order to follow the progression of meiotic prophase. The expression of *Setx* started to increase at 17 dpp and showed an additional striking increase at 31 dpp (**Figure 5A**), concurrent with the presence of spermatocytes and early spermatids. Expression levels decreased by 8 weeks, which may reflect the abundance of maturing spermatids in the testis germ-cell population. The lack of *Setx* expression in germ-cell deficient testes of *Kit^W/W-v^* and *Kit^Sl^* adult mice (**Figure 5A**) indicates indicated that *Setx* expression is restricted mainly to germ cells and not somatic cells of the testis. Further analysis of published scRNA-seq data of WT testicular cells [28] confirmed expression of *Setx* primarily in meiotic spermatocytes, with relatively low expression in some Sertoli cells and spermatids (**Figure 5B**).

**Figure 5.**
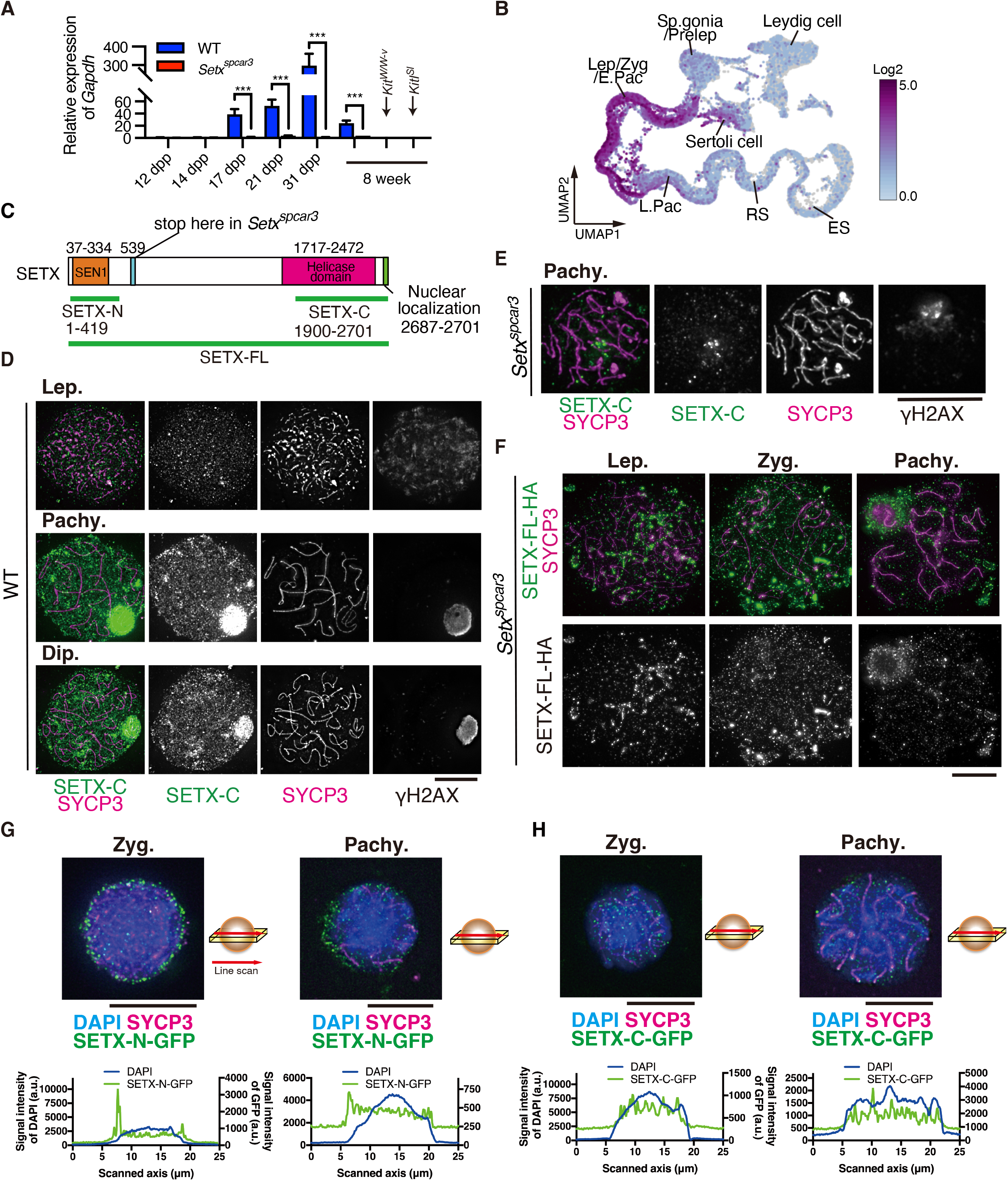
Expression of *Setx* and SETX in meiotic spermatocytes. **A.** Expression profile of *Setx* relative to *Gapdh* expression during the first wave of spermatogenesis by quantitative RT-PCR. Blue and red columns represent WT and *Setx^spcar3^* testis. Bars indicate standard deviation. Statistical significance was examined by non-parametric one-way ANOVA followed by Turkey’s test. *** *P* < 0.005. **B.** Uniform manifold approximation and projection (UMAP) representation for *Setx* expression in the testicular cells using multiplexed scRNA-seq data from 5-25 dpp WT mouse testis [28]. Color gradient represents the expression level of *Setx* in a log2 of arbitrary unit. Sp.gonia: spermatogonia. Prelep: preleptotene spermatocytes. Lep/Zyg: leptotene and zygotene spermatocytes. Pac.: pachytene spermatocyte. RS: round spermatid. ES: elongating spermatid. **C**. Domain structure of SETX protein. Numbers above the box represent regions of domains in amino acids. The green bar represents the region used as an antigen for the rabbit anti-SETX antibody. **D-E**. Localization of SETX in spermatocytes. Immunolabeling spread chromatin of spermatocytes from WT mice showed stage-specific localization of endogenous SETX in the autosome and the XY chromosomes (D). Chromatin localization of SETX was lost in the *Setx^spcar3^* spermatocytes (E). Endogenous SETX is shown in green. SYCP3 (magenta) is immunolabeled for staging, and γH2AX (grey) is the phosphorylated histone H2AFX formed near DNA damage. **F.** Transient expression of the HA-tagged full-length SETX (SETX-FL-HA) in the *Setx^spcar3^* spermatocytes after *in vivo* testis electroporation. SETX-FL-HA is shown in green. SYCP3 (magenta) is immunolabeled for staging. **G-H.** Transient expression of the GFP-tagged N- and C-terminus of SETX (SETX-N-GFP and SETX-C-GFP) in the WT spermatocytes after *in vivo* testis electroporation. Truncated SETX is shown in green. SYCP3 (magenta) is immunolabeled for staging. DAPI (blue) labels DNA. Sections of meiocytes are shown. Line plots represent the cellular distribution of truncated SETX (green) and DNA (blue). Scale bars are 10 µm.

We then examined the subnuclear localization of SETX in spread chromatin of spermatocytes using a polyclonal antibody against the C-terminus helicase domain of mouse SETX (aa. 1900-2701, **Figure 5C**). During early prophase, SETX exhibited a punctate localization pattern on the chromatin of WT leptotene spermatocytes (**Figure 5D**). During later prophase, SETX was found in punctate localization in the autosomal chromatin of WT pachytene or diplotene spermatocytes, as well as accumulated in the sex body (**Figure 5D**), confirming previous observations [22, 43]. These immuno-signals were greatly diminished in the *Setx^spcar3^* spermatocytes (**Figure 5E**).

We used transient expression assays to further assess nuclear localization of discrete domains of SETX, specifically truncated C- and N-terminus and full-length SETX (SETX-N-GFP, SETX-C-GFP, and SETX-FL-HA, respectively). Expression was achieved after *in vivo* testis electroporation (tEP) (see Methods) [31]. The localization pattern of transiently expressed SETX-FL-HA in the SETX-deficient *Setx^spcar3^*spermatocytes confirmed that of antibody detecting the endogenous SETX (**Figure 5F**). In addition, the SETX-N-GFP (aa. 1-419) construct was designed to cover the conserved sen1 domain, similar to the predicted *Setx^spcar3^* truncated SETX (**Figure 5C**). Because the SETX-N-GFP lacks a nucleotide-binding helicase domain and a nuclear localization signal (NLS), the majority of transiently expressed SETX-N in the WT meiocytes was observed outside of the DAPI-positive region (**Figure 5G**), indicating a lack of the ability to localize within the nucleus. On the other hand, localization of SETX-C-GFP (aa. 1900-2701), containing a major part of the helicase domain and a nuclear localization signal, was similar to the nuclear localization of SETX in WT meiocytes. However, localization of SETX-C-GFP in the XY sex chromatin was not observed in pachytene spermatocytes (**Figure 5H**), indicating that the sen1 domain is required for XY localization.

### *The loss of* Setx *caused the accumulation of unresolved R-loop in the* Setx^spcar3^ *testicular germ cells*

Given the role of SETX in resolving R-loop structures [22, 44, 45], unresolved R-loops could mark sites of DNA damage in the spermatocytes absent of SETX. We assessed R-loop formation in the testis from WT and *spcar3* mice using the S9.6 antibody, which recognizes DNA/RNA hybrids [46]. As expected, control spermatocytes did not show obvious R-loop accumulation in either RNaseH-treated and -untreated conditions (**Figure 6A**). However, in the testes of *Setx^spcar3^*mice, accumulation of R-loops was observed in meiotic spermatocytes and spermatogonia, together with non-specific signals in the cytoplasm of *Setx^spcar3^*cells (**Figure 6B**). The R-loop signals in the nucleus of *Setx^spcar3^* cells were diminished when treated with RNaseH (**Figure 6B**), confirming the specificity of S9.6 antibody. These results suggest that the loss of senataxin led to the accumulation of unresolved R-loops in germ cells of *Setx^spcar3^*males.

**Figure 6.**
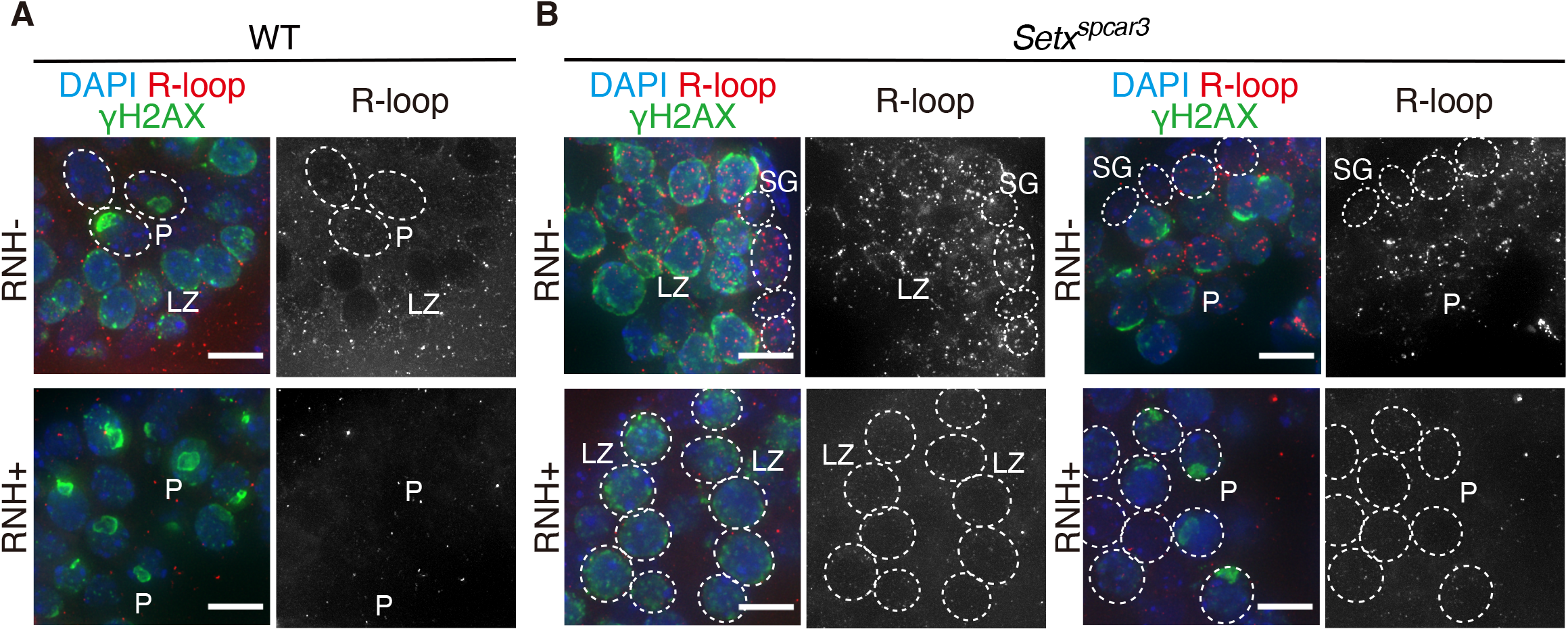
Loss of SETX causes accumulation of unresolved R-loops. **A-B**. Abnormal accumulation of unresolved R-loop in the *Setx^spcar3^* testicular cells. Few R-loop are detected either with or without RNaseH treatment (RNH+ or RNH-) in the meiotic spermatocytes of the WT mice (testicular section) using the 9.6 antibody (A). Testicular germ cells of the *Setx^spcar3^* mouse exhibit accumulation of R-loop signals without RNaseH treatment. The R-loop signals disappear after RNaseH treatment (B). LZ: leptotene and zygotene spermatocytes. P: pachytene spermatocytes. SG: spermatogonia. RNH: RNaseH treatment. DNA was stained using DAPI. Scale bars are 10 µm.

## DISCUSSION

### *A two-bp deletion in Setx gene, encoding a DNA/RNA helicase, in* Setx^spcar3^ *mutants*

Phenotype screening after ENU mutagenesis [20] identified the male-infertile *Setx^spcar3^* mutant phenotype. Linkage mapping and deep sequencing revealed a two-nucleotide deletion in the *Setx* gene. Because ENU usually causes one-nucleotide substitutions rather than deletions, it is possible that the *Setx^spcar3^* mutation might be spontaneous rather than mutagen-induced . The *Setx^spcar3^* mutation created a premature stop codon, resulting in a loss of *Setx* transcript in the testis of the *Setx^spcar3^* mutant mouse. After the genotype-confirmed backcross mating over 30 generations, we confirmed that the C3H congenic *Setx^spcar3^* homozygous mice lacked SETX protein and exhibited the *spcar3* phenotype, and, moreover, the histological and cytological phenotypes of *Setx^spcar3^* resembled those of *Setx*-KO mice [47]. Interestingly, our evaluation of fertility revealed that the female *Setx^spcar3^* homozygous mice exhibited a reduction in their litter size. This implies a potential role of SETX in female, as well as male, fertility [21, 22].

### *Spermatogenic and meiotic defects in the* Setx^spcar3^ *mice*

The *Setx^spcar3^* and *Setx*-KO mice [22] both exhibit arrest of spermatogenesis during the pachytene stage of meiotic prophase I. Our cytological analyses revealed that while *Setx^spcar3^* pachytene spermatocytes with fully synapsed SCs were frequently observed, no *Setx^spcar3^* spermatocytes progressed to the later pachytene stages marked by accumulation of histone H1t, suggesting that the spermatogenesis of *Setx^spcar3^*mice is arrested during at an early to mid-pachytene stage, after chromosome pairing but prior to recombination repair-based exchange. Because homologous pairing is mediated primarily by processes by which SPO11/TOPO6BL complex-programmed DSBs are repaired through homologous recombination [5, 48], aberrant formation of DSBs could result in asynapsis [49-52]. Consistent with their phenotype, the *Setx^spcar3^* pachytene spermatocytes exhibited unrepaired DSBs, marked by γH2AX and DNA damage repair factors (DMC1, RAD51, and SPATA22) remaining on the fully synapsed AEs of the autosomes, while MLH1, marking exchange, was not detected.

### *Defective resolution of R-loop in the testicular cells in* Setx^spcar3^ *mice*

Mammalian SETX is a putative DNA/RNA helicase [44, 53] and its yeast ortholog sen1 has unwinding activity for DNA/RNA hybrids [54-56]. Mutations in sen1 cause transcription-associated DNA damage [45, 57] that is presumably due to unresolved R-loop structure. Both formation and resolution of R-loops are critical for gene transcription directly [44, 58-60] or indirectly [61, 62]. A loss of SETX function leads to the accumulation of DNA damage in human cells [63] and mouse spermatocytes [22]. The *Setx^spcar3^* phenotype highlights the indispensable role of mammalian SETX in genome integrity during meiosis, and suggests the importance of R-loop unwinding function. Specifically, pachytene spermatocytes from both *Setx*-KO mouse [22] and *Setx^spcar3^* mouse (present study) exhibited an accumulation of unresolved R-loops. Our observations reveal R-loop accumulation not only in *Setx^spcar3^*spermatocytes but also in Leydig cells and Sertoli cells, which disappeared when treated with RNaseH. These observations suggested an essential and ubiquitous role of SETX in the resolution of R-loop not only in pachytene spermatocytes but possibly also in somatic testicular cells.

Our cytological analyses of SETX in meiocytes revealed endogenous SETX was in the nucleus of spermatocytes throughout meiotic prophase I. Interestingly, SETX exhibited substantial accumulation at the XY chromosomes of pachytene spermatocytes (Figure 5) [22]. It is not known if R-loops are formed in the transcriptionally inactive XY chromosomes during pachytene stage. Given that *Setx^spcar3^*pachytene spermatocytes exhibited defective formation of the XY body, SETX may have an independent role in the XY chromosomes, perhaps in MSCI [22, 43]. Our transient expression analyses revealed that SETX-C-GFP, which lacks the sen1 domain, did not show accumulation in the XY chromosomes. This may indicate that the sen1 domain is required for the XY localization of SETX during the pachytene stage; however, the DNA/RNA-binding helicase domain is also indispensable. Furthermore, our study showed that ectopic SETX-FL in *Setx^spcar3^* pachytene spermatocytes colocalized with γH2AX and weakly with the AEs, suggesting that the ectopic SETX was recruited to the DNA damage sites or/and R-loop sites even in *spcar3* pachytene spermatocytes.

Taken together, because SETX is known to be essential for the resolution of harmful R-loops, it is plausible that unresolved R-loops caused the persistent autosomal DNA DSBs in *Setx^spcar3^* pachytene spermatocytes. Whether DNA DSBs observed in the *Setx^spcar3^* pachytene spermatocytes were associated with transcription-derived R-loops is still not clear, and further studies with new mouse models or assays will be required to resolve this.

### Setx *models for disease*

These results establish the *Setx^spcar3^* mutant mouse as a model for male infertility, especially given that AOA2 patients carrying loss-of-function mutations in the *SETX* gene exhibit spermatogenesis defects [64]. Additionally, our phenotypic analyses also revealed reduced fertility in female *Setx^spcar3^*mice, as also reported for the female *Setx*-KO mice [21, 22]. Like *Setx*-defective spermatocytes, aged female *Setx*-KO oocytes exhibited unrepaired DMA damage [63]. Such DNA damage could be a source of detrimental mutations, increasing the risk of heritable genetic diseases for the next generations. Therefore, the mouse *Setx* mutations provide important models for investigating infertility and genome integrity in both male and female mice.

In humans, mutations in SETX cause neurodegenerative diseases, such as amyotrophic lateral sclerosis type 4 (ALS4) [64-67] and ataxia oculomotor apraxia type 2 (AOA2) [68, 69], by gain-of-function and loss-of-function, respectively. A loss of SETX function leads to RNA degradation and ultimately degeneration of motor neurons [63, 70], probably through either or combination of genome instability, abnormal DNA/histone modifications, and transcription by affecting regulatory factors [71]. In contrast, the *spcar3* mice and *Setx*-KO mice, at least at young ages, exhibit no apparent neuronal dysfunction in addition to the male-restricted infertility [22]. Possibly SETX-associated neurodegenerative diseases in humans may not be caused by R-loop accumulation, and indeed, R-loop accumulation was not observed in the brains of the *Setx*-KO mice at young ages [72]. Therefore, further studies are required to elucidate the pathogenic mechanisms of neurological defects associated with SETX deficiency [73, 74].

In summary, this study identifies a new ENU-induced allele of the mouse *Setx* gene, *Setx^spcar3^*, on a defined genetic background. Similar to a previously reported knockout allele, this is also associated with meiotic failure and male fertility and will provide a new resource for further study of senataxin structure and the role of R-loops during meiosis and gametogenesis. Such “forward genetic” screens (identifying gene from induced phenotype) are likely to continue to be productive in the current genome sequencing era. Moreover, induced base pair-specific alleles, such as *Setx^spcar3^*, with a potentially “fixable” small lesion, are good models for gene-editing therapies that are on the horizon.

## AUTHOR CONTRIBUTION

Y.F., Y.O., and M.A.H. conceived the project. Experiments were carried out by Y.F., K.S., F.S., S.P., and E.I.. J.S. and M.A.H generated the *spcar3* mouse. Y.F., Y.O., and M.A.H wrote the paper.

## ACKNOWLEDGEMENTS

We thank Handel and Okada laboratory members for their comments and discussion. We thank Lucy Rowe and Mary Barter (The Jackson Laboratory, Bar Harbor, USA) for the preliminary mapping analysis. We acknowledge the Scientific Services of The Jackson Laboratory for outstanding support. We thank Dr. Tetsuo Kunieda for the travel support to YF. We thank Drs. Kei-ichiro Ishiguro, Haruhiko Siomi, and Bernard de Massy for helpful discussion.

## REFERENCES

1. Baudat F, Manova K, Yuen JP, Jasin M, Keeney S. Chromosome synapsis defects and sexually dimorphic meiotic progression in mice lacking *Spo11*. Mol Cell 2000; 6:989–998.

2. Romanienko PJ, Camerini-Otero RD. The mouse *Spo11* gene is required for meiotic chromosome synapsis. Mol Cell 2000; 6:975–987.

3. Robert T, Nore A, Brun C, Maffre C, Crimi B, Bourbon HM, de Massy B. The TopoVIB-Like protein family is required for meiotic DNA double-strand break formation. Science 2016; 351:943–949.

4. Vrielynck N, Chambon A, Vezon D, Pereira L, Chelysheva L, De Muyt A, Mezard C, Mayer C, Grelon M. A DNA topoisomerase VI-like complex initiates meiotic recombination. Science 2016; 351:939–943.

5. Bhalla N, Dernburg AF. Prelude to a division. Annu Rev Cell Dev Biol 2008; 24:397–424.

6. Barzel A, Kupiec M. Finding a match: how do homologous sequences get together for recombination? Nat Rev Genet 2008; 9:27–37.

7. Keeney S, Lange J, Mohibullah N. Self-organization of meiotic recombination initiation: general principles and molecular pathways. Annu Rev Genet 2014; 48:187–214.

8. Cahoon CK, Hawley RS. Regulating the construction and demolition of the synaptonemal complex. Nat Struct Mol Biol 2016; 23:369–377.

9. Fujiwara Y, Horisawa-Takada Y, Inoue E, Tani N, Shibuya H, Fujimura S, Kariyazono R, Sakata T, Ohta K, Araki K, Okada Y, Ishiguro KI. Meiotic cohesins mediate initial loading of HORMAD1 to the chromosomes and coordinate SC formation during meiotic prophase. PLoS Genet 2020; 16:e1009048.

10. Ishiguro K, Kim J, Shibuya H, Hernandez-Hernandez A, Suzuki A, Fukagawa T, Shioi G, Kiyonari H, Li XC, Schimenti J, Hoog C, Watanabe Y. Meiosis-specific cohesin mediates homolog recognition in mouse spermatocytes. Genes Dev 2014; 28:594–607.

11. 11. Fujiwara Y. Differential gene expression revealed by transcriptomic analyses of male germ cells. J Anim Genet 2014; 42:91-99.

12. Kota SK, Feil R. Epigenetic transitions in germ cell development and meiosis. Dev Cell 2010; 19:675–686.

13. Fallahi M, Getun IV, Wu ZK, Bois PRJ. A global expression switch marks pachytene initiation during mouse male meiosis. Genes 2010; 1:469–483.

14. Shima JE, McLean DJ, McCarrey JR, Griswold MD. The murine testicular transcriptome: characterizing gene expression in the testis during the progression of spermatogenesis. Biol Reprod 2004; 71:319–330.

15. Handel MA. The XY body: a specialized meiotic chromatin domain. Exp Cell Res 2004; 296:57–63.

16. Turner JM. Meiotic sex chromosome inactivation. Development 2007; 134:1823–1831.

17. Handel MA. The XY body: an attractive chromatin domain. Biol Reprod 2020; 102:985–987.

18. Alavattam KG, Maezawa S, Andreassen PR, Namekawa SH. Meiotic sex chromosome inactivation and the XY body: a phase separation hypothesis. Cell Mol Life Sci 2021; 79:18.

19. Schimenti JC, Handel MA. Unpackaging the genetics of mammalian fertility: strategies to identify the "reproductive genome". Biol Reprod 2018; 99:1119–1128.

20. Handel MA, Lessard C, Reinholdt L, Schimenti J, Eppig JJ. Mutagenesis as an unbiased approach to identify novel contraceptive targets. Mol Cell Endocrinol 2006; 250:201–205.

21. Subramanian GN, Lavin M, Homer HA. Premature ovarian ageing following heterozygous loss of Senataxin. Mol Hum Reprod 2021; 27.

22. Becherel OJ, Yeo AJ, Stellati A, Heng EY, Luff J, Suraweera AM, Woods R, Fleming J, Carrie D, McKinney K, Xu X, Deng C, et al. Senataxin plays an essential role with DNA damage response proteins in meiotic recombination and gene silencing. PLoS Genet 2013; 9:e1003435.

23. Bolger AM, Lohse M, Usadel B. Trimmomatic: a flexible trimmer for Illumina sequence data. Bioinformatics 2014; 30:2114–2120.

24. Andrews S. FastQC: A Quality Control Tool for High Throughput Sequence Data. In; 2010.

25. Dobin A, Davis CA, Schlesinger F, Drenkow J, Zaleski C, Jha S, Batut P, Chaisson M, Gingeras TR. STAR: ultrafast universal RNA-seq aligner. Bioinformatics 2013; 29:15–21.

26. Li H, Handsaker B, Wysoker A, Fennell T, Ruan J, Homer N, Marth G, Abecasis G, Durbin R, Genome Project Data Processing S. The Sequence Alignment/Map format and SAMtools. Bioinformatics 2009; 25:2078–2079.

27. Li B, Dewey CN. RSEM: accurate transcript quantification from RNA-Seq data with or without a reference genome. BMC Bioinformatics 2011; 12:323.

28. Ernst C, Eling N, Martinez-Jimenez CP, Marioni JC, Odom DT. Staged developmental mapping and X chromosome transcriptional dynamics during mouse spermatogenesis. Nat Commun 2019; 10:1251.

29. Zheng GX, Terry JM, Belgrader P, Ryvkin P, Bent ZW, Wilson R, Ziraldo SB, Wheeler TD, McDermott GP, Zhu J, Gregory MT, Shuga J, et al. Massively parallel digital transcriptional profiling of single cells. Nat Commun 2017; 8:14049.

30. He S, Gillies JP, Zang JL, Cordoba-Beldad CM, Yamamoto I, Fujiwara Y, Grantham J, DeSantis ME, Shibuya H. Distinct dynein complexes defined by DYNLRB1 and DYNLRB2 regulate mitotic and male meiotic spindle bipolarity. Nat Commun 2023; 14:1715.

31. 31. Shibuya H, Morimoto A, Watanabe Y. The Dissection of Meiotic Chromosome Movement in Mice Using an In Vivo Electroporation Technique. Plos Genetics 2014; 10.

32. Pendlebury DF, Fujiwara Y, Tesmer VM, Smith EM, Shibuya H, Watanabe Y, Nandakumar J. Dissecting the telomere-inner nuclear membrane interface formed in meiosis. Nat Struct Mol Biol 2017; 24:1064–1072.

33. Cobb J, Reddy RK, Park C, Handel MA. Analysis of expression and function of topoisomerase I and II during meiosis in male mice. Mol Reprod Dev 1997; 46:489–498.

34. Cobb J, Reddy RK, Park C, Handel MA. Analysis of expression and function of topoisomerase I and II during meiosis in male mice. Molecular Reproduction and Development 1999; 46:489–498.

35. Sun F, Handel MA. Regulation of the meiotic prophase I to metaphase I transition in mouse spermatocytes. Chromosoma 2008; 117:471–485.

36. La Salle S, Sun F, Zhang XD, Matunis MJ, Handel MA. Developmental control of sumoylation pathway proteins in mouse male germ cells. Dev Biol 2008; 321:227–237.

37. Larionov A, Krause A, Miller W. A standard curve based method for relative real time PCR data processing. BMC Bioinformatics 2005; 6:62.

38. Gentalen E, White T, Proctor J. Peggy™: size- or charge-based western blotting at the push of a button. Nature Methods 2013; 10:i-ii.

39. Cobb J, Cargile B, Handel MA. Acquisition of competence to condense metaphase I chromosomes during spermatogenesis. Dev Biol 1999; 205:49–64.

40. Handel MA, Schimenti JC. Genetics of mammalian meiosis: regulation, dynamics and impact on fertility. Nat Rev Genet 2010; 11:124–136.

41. Zhang J, Nandakumar J, Shibuya H. BRCA2 in mammalian meiosis. Trends Cell Biol 2022; 32:281–284.

42. Santucci-Darmanin S, Walpita D, Lespinasse F, Desnuelle C, Ashley T, Paquis-Flucklinger V. MSH4 acts in conjunction with MLH1 during mammalian meiosis. FASEB J 2000; 14:1539–1547.

43. Yeo AJ, Becherel OJ, Luff JE, Graham ME, Richard D, Lavin MF. Senataxin controls meiotic silencing through ATR activation and chromatin remodeling. Cell Discovery 2015; 1:15025.

44. Skourti-Stathaki K, Proudfoot NJ, Gromak N. Human senataxin resolves RNA/DNA hybrids formed at transcriptional pause sites to promote Xrn2-dependent termination. Mol Cell 2011; 42:794–805.

45. Mischo HE, Gomez-Gonzalez B, Grzechnik P, Rondon AG, Wei W, Steinmetz L, Aguilera A, Proudfoot NJ. Yeast Sen1 helicase protects the genome from transcription-associated instability. Mol Cell 2011; 41:21–32.

46. Boguslawski SJ, Smith DE, Michalak MA, Mickelson KE, Yehle CO, Patterson WL, Carrico RJ. Characterization of monoclonal antibody to DNA.RNA and its application to immunodetection of hybrids. J Immunol Methods 1986; 89:123–130.

47. Bayes JJ, Malik HS. Altered heterochromatin binding by a hybrid sterility protein in *Drosophila* sibling species. Science 2009; 326:1538–1541.

48. Zickler D, Kleckner N. Recombination, Pairing, and Synapsis of Homologs during Meiosis. Cold Spring Harb Perspect Biol 2015; 7.

49. Pittman DL, Cobb J, Schimenti KJ, Wilson LA, Cooper DM, Brignull E, Handel MA, Schimenti JC. Meiotic prophase arrest with failure of chromosome synapsis in mice deficient for Dmc1, a germline-specific RecA homolog. Mol Cell 1998; 1:697–705.

50. Zhang J, Fujiwara Y, Yamamoto S, Shibuya H. A meiosis-specific BRCA2 binding protein recruits recombinases to DNA double-strand breaks to ensure homologous recombination. Nat Commun 2019; 10:722.

51. Zhang J, Gurusaran M, Fujiwara Y, Zhang K, Echbarthi M, Vorontsov E, Guo R, Pendlebury DF, Alam I, Livera G, Emmanuelle M, Wang PJ, et al. The BRCA2-MEILB2-BRME1 complex governs meiotic recombination and impairs the mitotic BRCA2-RAD51 function in cancer cells. Nat Commun 2020; 11:2055.

52. La Salle S, Palmer K, O’Brien M, Schimenti JC, Eppig J, Handel MA. *Spata22*, a novel vertebrate-specific gene, is required for meiotic progress in mouse germ cells. Biol Reprod 2012; 86:45.

53. Bennett CL, La Spada AR. Senataxin, A Novel Helicase at the Interface of RNA Transcriptome Regulation and Neurobiology: From Normal Function to Pathological Roles in Motor Neuron Disease and Cerebellar Degeneration. Adv Neurobiol 2018; 20:265–281.

54. Kim HD, Choe J, Seo YS. The sen1(+) gene of Schizosaccharomyces pombe, a homologue of budding yeast SEN1, encodes an RNA and DNA helicase. Biochemistry 1999; 38:14697–14710.

55. Leonaite B, Han Z, Basquin J, Bonneau F, Libri D, Porrua O, Conti E. Sen1 has unique structural features grafted on the architecture of the Upf1-like helicase family. EMBO J 2017; 36:1590–1604.

56. Martin-Tumasz S, Brow DA. Saccharomyces cerevisiae Sen1 Helicase Domain Exhibits 5’-to 3’-Helicase Activity with a Preference for Translocation on DNA Rather than RNA. J Biol Chem 2015; 290:22880–22889.

57. Alzu A, Bermejo R, Begnis M, Lucca C, Piccini D, Carotenuto W, Saponaro M, Brambati A, Cocito A, Foiani M, Liberi G. Senataxin associates with replication forks to protect fork integrity across RNA-polymerase-II-transcribed genes. Cell 2012; 151:835–846.

58. Ginno PA, Lott PL, Christensen HC, Korf I, Chedin F. R-loop formation is a distinctive characteristic of unmethylated human CpG island promoters. Mol Cell 2012; 45:814–825.

59. Nakama M, Kawakami K, Kajitani T, Urano T, Murakami Y. DNA-RNA hybrid formation mediates RNAi-directed heterochromatin formation. Genes Cells 2012; 17:218–233.

60. Castellano-Pozo M, Santos-Pereira JM, Rondon AG, Barroso S, Andujar E, Perez-Alegre M, Garcia-Muse T, Aguilera A. R loops are linked to histone H3 S10 phosphorylation and chromatin condensation. Mol Cell 2013; 52:583–590.

61. Skourti-Stathaki K, Torlai Triglia E, Warburton M, Voigt P, Bird A, Pombo A. R-Loops Enhance Polycomb Repression at a Subset of Developmental Regulator Genes. Mol Cell 2019; 73:930–945 e934.

62. Grunseich C, Wang IX, Watts JA, Burdick JT, Guber RD, Zhu Z, Bruzel A, Lanman T, Chen K, Schindler AB, Edwards N, Ray-Chaudhury A, et al. Senataxin Mutation Reveals How R-Loops Promote Transcription by Blocking DNA Methylation at Gene Promoters. Mol Cell 2018; 69:426–437 e427.

63. Suraweera A, Becherel OJ, Chen P, Rundle N, Woods R, Nakamura J, Gatei M, Criscuolo C, Filla A, Chessa L, Fusser M, Epe B, et al. Senataxin, defective in ataxia oculomotor apraxia type 2, is involved in the defense against oxidative DNA damage. J Cell Biol 2007; 177:969–979.

64. Chen YZ, Bennett CL, Huynh HM, Blair IP, Puls I, Irobi J, Dierick I, Abel A, Kennerson ML, Rabin BA, Nicholson GA, Auer-Grumbach M, et al. DNA/RNA helicase gene mutations in a form of juvenile amyotrophic lateral sclerosis (ALS4). Am J Hum Genet 2004; 74:1128–1135.

65. 65. Tripolszki K, Torok D, Goudenege D, Farkas K, Sulak A, Torok N, Engelhardt JI, Klivenyi P, Procaccio V, Nagy N, Szell M. High-throughput sequencing revealed a novel SETX mutation in a Hungarian patient with amyotrophic lateral sclerosis. Brain Behav 2017; 7:e00669.

66. Grunseich C, Patankar A, Amaya J, Watts JA, Li D, Ramirez P, Schindler AB, Fischbeck KH, Cheung VG. Clinical and Molecular Aspects of Senataxin Mutations in Amyotrophic Lateral Sclerosis 4. Ann Neurol 2020; 87:547–555.

67. Groh M, Albulescu LO, Cristini A, Gromak N. Senataxin: Genome Guardian at the Interface of Transcription and Neurodegeneration. J Mol Biol 2017; 429:3181–3195.

68. Fogel BL, Lee JY, Perlman S. Aberrant splicing of the senataxin gene in a patient with ataxia with oculomotor apraxia type 2. Cerebellum 2009; 8:448–453.

69. Arning L, Schols L, Cin H, Souquet M, Epplen JT, Timmann D. Identification and characterisation of a large senataxin (SETX) gene duplication in ataxia with ocular apraxia type 2 (AOA2). Neurogenetics 2008; 9:295–299.

70. Chen YZ, Hashemi SH, Anderson SK, Huang Y, Moreira MC, Lynch DR, Glass IA, Chance PF, Bennett CL. Senataxin, the yeast Sen1p orthologue: characterization of a unique protein in which recessive mutations cause ataxia and dominant mutations cause motor neuron disease. Neurobiol Dis 2006; 23:97–108.

71. Groh M, Gromak N. Out of balance: R-loops in human disease. PLoS Genet 2014; 10:e1004630.

72. Yeo AJ, Becherel OJ, Luff JE, Cullen JK, Wongsurawat T, Jenjaroenpun P, Kuznetsov VA, McKinnon PJ, Lavin MF. R-loops in proliferating cells but not in the brain: implications for AOA2 and other autosomal recessive ataxias. PLoS One 2014; 9:e90219.

73. Catford SR, O’Bryan MK, McLachlan RI, Delatycki MB, Rombauts L. Germ cell arrest associated with a SETX mutation in ataxia oculomotor apraxia type 2. Reprod Biomed Online 2019; 38:961–965.

74. Becherel OJ, Fogel BL, Zeitlin SI, Samaratunga H, Greaney J, Homer H, Lavin MF. Disruption of Spermatogenesis and Infertility in Ataxia with Oculomotor Apraxia Type 2 (AOA2). Cerebellum 2019; 18:448–456.

